# Extra-genital wounding delays remating in the sexually cannibalistic springbok mantis

**DOI:** 10.1101/2025.08.21.671560

**Authors:** Nathan W. Burke, Laura Knapwerth, Jutta M. Schneider

## Abstract

A fascinating consequence of sexual conflict in animals is the maintenance of traits in males that cause physical damage to females during mating interactions. Such harm is hypothesised to be either an adaptation that enhances male fitness, or a collateral side-effect of adaptations that benefit males in other contexts. Tests of these hypotheses have mostly focused on traumatic copulation, where males wound females internally with weaponised genitalia, despite the widespread occurrence of extra-genital injury inflicted by non-genitalic structures like teeth, fangs or claws. Here, we take advantage of the unique mating interactions of the sexually cannibalistic springbok mantis, *Miomantis caffra*, to investigate the evolution of extra-genital wounding. Males of this species use their foretibial claws to stab females in the abdomen while fighting back against cannibalistic attacks during mating attempts. If stabbing females is adaptive, we predicted that experimentally wounded females would alter their remating behaviour or reproductive scheduling to the benefit of their mates. We found that injured females did not differ in their attractiveness or remating likelihood compared to intact females, and did not show an enhanced propensity to attack second courting males. Injured females also showed no change in mortality, fecundity, or offspring production that would suggest terminal investment due to male manipulation through wounding. However, our experiments revealed a statistically significant delay in the timing of remating among injured females, which could be interpreted as a benefit to males if such a delay reduces sperm competition. But given that remating was not prevented, only deferred, the injury-induced delay we observed is unlikely to be ecologically important, although field studies would be required to confirm this. Taken together, our results suggest that extra-genital wounding in the springbok mantis is unlikely to be adaptive but may instead be a pleiotropic side-effect of males trying to avoid being cannibalised.

## INTRODUCTION

Sexual conflict is ubiquitous in animals and occurs when the evolutionary interests of males and females are different and cannot be achieved at the same time (Arnqvist & Rowe, 2005; Parker, 1979). When the sexes are in conflict over mating outcomes, selection is expected to act on males and females in opposing directions, leading to the evolution of antagonistic traits that help each sex to optimise their own fitness at the expense of their mating partner, such as grasping structures in males and anti-grasping structures in females (Arnqvist & Rowe, 2002; Chapman, 2006; Chapman et al., 2003; Parker, 1979; Perry & Rowe, 2015). Such sexually antagonistic selection can result in diverse outcomes, including runaway arms races between coercion and resistance, stable compromises between the sexes, and female indifference (Gavrilets, 2000; Härdling & Smith, 2005; Holland & Rice, 1998; Rowe et al., 2005).

Numerous examples of antagonistic traits in males involve harm to females. In some species, males produce toxic substances in their ejaculates or pheromones that alter female reproductive decisions (Chapman et al., 1995; Fricke et al., 2009; Kimura & Chiba, 2015; Moore et al., 2003). In others, males traumatically mate with females using weaponised genitalia that cause internal injury, reducing lifespan and fitness (Blanckenhorn et al., 2002; Crudgington & Siva-Jothy, 2000; Morrow & Arnqvist, 2003). The evolution of such male-inflicted harm has been hypothesised to occur via two mechanisms. The most parsimonious explanation is that harm is an incidental side-effect of selection acting on other traits that otherwise benefit males, such as structures that facilitate internal or external anchorage to females during copulation (Parker, 1979). According to this ‘pleiotropic harm hypothesis’, injury is maintained by selection not because it directly benefits males, but because the cost of injury in terms of reduced reproductive output of injured mates does not outweigh the benefit that males gain by using traits that result in harm (Parker, 1979). The second idea is that male-inflicted injury is favoured because it directly benefits male fitness. This ‘adaptive harm hypothesis’ posits that harm is not incidental but a consequence of selection acting on males to selfishly alter either future female mating behaviour or reproductive scheduling (Johnstone & Keller, 2000; Lessells, 2005). In the first case, injury is adaptive if it causes females to decrease the rate at which they remate with other males, thereby reducing the potential for sperm competition (Johnstone & Keller, 2000).

Alternatively, injury is adaptive if it triggers females into a terminal investment response whereby resources are immediately redirected to present reproduction at the expense of future reproduction because of the negative effect that injury has on female survival (Lessells, 2005; Michiels, 1998). The current consensus is that most examples of male-inflicted injury are pleiotropic, arising as an essentially unavoidable by-product of male traits that are adaptive in other contexts, such as in sperm competition or male-male combat (e.g., see Edvardsson & Tregenza, 2005; Hotzy & Arnqvist, 2009; Morrow et al., 2003; Tong et al., 2021). However, almost all tests of male-imposed harm have been performed on species where injury is inflicted internally to the female reproductive tract by weaponised genitalia, even though males are known to harm females in a multitude of other ways during mating interactions (e.g., see Daly, 1978; Kimura & Chiba, 2015; Lange et al., 2013; Le Boeuf & Mesnick, 1990; Pratt & Carrier, 2001).

External harm occurs in a number of sexually cannibalistic taxa where females attack and consume realised or potential mating partners before, during, or after mating (Elgar, 1992). Notable examples include males using pedipalps to mutilate the external genitalia (scapi) of females in orb-weaving spiders (Mouginot & Uhl, 2019; Nakata, 2016) and males injecting venom through fang bites in crab spiders (Sentenská et al., 2020). Such behaviours are almost certainly the product of sexually antagonistic selection (Burke, 2024; Schneider, 2014), but little is known about how they affect fitness. External genital mutilation in orb-weavers apparently benefits injurious males by preventing remating (Mouginot et al., 2015), but whether male-inflicted external damage in other cannibalistic taxa is also adaptive remains unknown. An important distinction between males of cannibalistic versus non-cannibalistic taxa is that while both are selected to secure matings, males of cannibalistic species also typically try to avoid being eaten by females (Lelito & Brown, 2006; but see Andrade, 1996). This means that mating behaviours that harm females could be a side-effect of selection acting on males to avoid being cannibalised. On the other hand, consuming males is known to boost female fecundity (Barry et al., 2008; Knapwerth & Burke, 2025), and males could therefore benefit by causing harm if injured females increase attacks on subsequent suitors, thereby reducing the risk of sperm competition for injurious males.

Females of the springbok mantis, *Miomantis caffra*, are noncopulatory cannibals that attack and consume males exclusively without mating, and at very high rates (Knapwerth & Burke, 2024; Pollo et al., 2021). But males of this species have evolved an effective tactic to avoid being eaten. Rather than fleeing from cannibalistic attacks like males of other species (e.g., Barry et al., 2009; Lawrence, 1992), *M. caffra* males fight back against aggressive females in violent premating struggles that enhance the likelihood of mating and reduce the chance of cannibalism if males win the interaction (Burke & Holwell, 2021). However, during these stereotyped wrestling bouts, males often stab females in the abdomen with their foretibial claws, resulting in sometimes significant hemolymph loss (Burke & Holwell, 2023a) and even prolapse of the intestines through the wound (personal observation, NWB). Despite the severity of these injuries, not all struggles are harmful. Why some females are stabbed and not others is currently unknown, and the extent to which female lifespan and reproduction are affected by injury is also yet to be comprehensively investigated. It is therefore unclear whether males stab females deliberately to manipulate female life-history in their favour, or incidentally as a side effect of a drive to stay alive.

The type of external, extra-genital harm that occurs in *M. caffra* offers an unprecedented opportunity to use phenotypic engineering to simulate injury as it occurs in natural male-female interactions. To investigate whether injuring females provides benefits to males as per the adaptive harm hypothesis, we experimentally injured *M. caffra* females the same way that males stab female abdomens and observed female responses in terms of remating behaviour and reproductive scheduling. If injury is adaptive, we predicted that males would gain by causing injured females to be less open to remate with other males, resulting in reduced attractiveness in such females, lower likelihood to remate, delayed remating, increased likelihood of avoiding remating through noncopulatory cannibalism and/or increased likelihood of ovipositing without remating. If males adaptively induce a terminal investment response by injuring females, we predicted that injured females would die sooner than intact females, lay more of their eggs earlier than later, and produce more offspring in early life.

## METHODS

### Insect husbandry

Mantises were F2 individuals hatched from laboratory stocks originally established from oothecae collected from several parks and gardens in Auckland, New Zealand, where the species is an introduced pest. Mantises were raised in the laboratory under constant temperature (25^°^C) and humidity (60%), and a 10:14h day:night cycle. We housed mantises in their own separate upturned plastic cups that varied in volume according to developmental stage (50 mL: 1^st^ to 3^rd^ instar; 250 mL: 4^th^ to penultimate instar; 400 mL: adulthood). To facilitate ease of perching and moulting, the insides of the 50 mL and 250 mL cups were mechanically abraded and a hole was drilled in the ceiling and stuffed with dampened cottonwool, while the 400 mL cups had a ceiling of fine mesh and a strip of the same material attached to the wall. All mantises were fed live prey twice a week, with 1^st^ to 3^rd^ instars receiving several fruit flies (*Drosophila sp*. flightless hybrids), 3^rd^ to 5^th^ instars receiving 2 green bottle flies (*Lucilia sp*.), and 5^th^ to penultimate instars receiving 2 blow flies (*Calliphora sp*.). As adults, the sexes were fed differently due to sexual size dimorphism, with adult males receiving 2 green bottle flies and adult females 3 blow flies. Females were also occasionally fed medium-sized crickets (*Gryllus bimaculatus*). Cups were watered twice a week at feeding time for mantises to drink. The pronotum length of all individuals was measured with callipers at adulthood as a proxy for body size.

### Treatment allocation

To determine whether injury is adaptive for males, we conducted a laboratory experiment that assessed the reproductive consequences of males injuring females. To do this, we randomly allocated females to either an “injured” treatment group (*n* = 38) or an “intact” control (*n* = 36). Females in the injured group were temporarily anaesthetised with CO_2_ for 10 secs and stabbed twice in the abdomen in the same way that males typically injure females during premating struggles (i.e., on the ventral side of the abdomen, one wound to the left and one to the right, on the 4^th^ or 5^th^ sternite). Stabbing was achieved using a bespoke implement consisting of a dead males’ foretibial claw (which is the injurious part of the male’s foreleg) attached to the blunt end of a fine paintbrush. This injuring procedure ensured that the experimentally applied stab wounds were of similar size and depth to natural stab wounds. The stabbing implement was cleaned with ethanol between stabbings. Females in the intact treatment group were anaesthetised the same way as injured females but were not stabbed. We deliberately did not apply an injury procedural control to intact females (such as by injuring an inconsequential body part like the non-functional wings) because any form of injury could potentially trigger a behavioural or life-history response (see Morrow et al., 2003), and we wanted to avoid conflating injury effects. Injured and intact females were of similar age and size at treatment allocation (time since final moult: injured: 30.53 ± 1.14 days, intact: 29.72 ± 1.4 days; pronotum length: injured: 15.22 ± 0.11 mm, intact: 15.38 ± 0.15 mm).

Females were mated with males immediately prior to treatment allocation on the same day. Mating was facilitated by pairing each female with a randomly selected virgin male inside a clear plastic enclosure (14.5 × 11.5 × 7.1 cm) with the inside walls and ceiling lined with paper towel for easy perching. Due to unexpectedly low male numbers, some females were paired with previously mated males. Mating was determined by the presence of a conspicuous spermatophore (Burke & Holwell, 2023b). Females were fed 5 blow flies the day before pairing to ensure they were well satiated, and 3 blowflies twice a week thereafter until mating occurred. Cannibalism was an exclusion criterion. Consequently, 22 females that cannibalised were immediately excluded from the experiment and replaced with other individuals. Excluded females did not contribute to sample sizes. To prevent males from injuring females, their foretibial claws were ablated with surgical scissors at least 1 day prior to pairing. All ablation surgeries were performed while males were anaesthetised with CO_2_. Thus, our experimental setup simulated a scenario where females were either injured or not due to mating interactions.

### Female attractiveness

To ascertain whether males are selected to injure females to make them less attractive to other males, we performed a mate choice experiment on injured and intact females soon after treatment allocation. To do this, we placed a pair of injured and intact females (selected haphazardly) inside separate upturned stainless-steel mesh containers (8.5 × 12.5 × 8.5 cm) positioned against opposite walls inside a fully-enclosed glass terrarium, with the left or right position determined randomly. Mesh containers were used to allow pheromones, which female mantises produce to attract males (Maxwell et al., 2010), to diffuse passively through the terrarium. Once enclosed females were in position, we added a randomly selected virgin male to the centre of the terrarium, positioned on the ceiling. We then waited for a period of 3 hrs for males to choose one of the females, and recorded which female was chosen at half hour intervals. We considered a choice to have been made if a male moved to within 1 cm of one or other of the mesh containers. Three trials were performed at a time, over 11 days. Mesh containers and terraria glass were cleaned with soapy water, dried with paper towel, then sanitised with 70% ethanol in between trails to ensure that any remnant pheromones or contact chemicals were thoroughly removed. Trials were conducted under red light, which mantises cannot see (Sontag, 1971), to simulate night-time conditions when males are typically most active in search for mates. We performed a total of 50 trials, 16 of which resulted in the male clearly choosing one or other of the females. A number of females from trials ending in no choice were reused in subsequent trials.

If injury alters female attractiveness to the benefit of injurious males, then we predicted that injured females would attract fewer males compared to intact females. To assess this prediction statistically, we performed an exact binomial test on the proportion of trials that ended in the male choosing the injured female. To additionally determine whether female attractiveness was correlated with male or female traits, we fitted a generalised linear model (GLM) with a binomial error structure in which the female that the male chose was the binary response variable (injured = 1, intact = 0). The numeric covariates in the model were male age (time from final moult to trial date, in days) and body size (pronotum length, in mm), and the difference in age and body size between female pairs (intact minus injured). Potential time effects and cage effects could not be accounted for due to model non-convergence when we attempted to include trial date (block) and cage ID in the model. We were also unable to account for potential relatedness effects since the fitting of terms for family identities as random effects resulted in model singularity.

### Remating and cannibalism

To test whether injury reduces female remating or increases cannibalism to the benefit of male paternity, we performed another mating experiment using new males and the same injured and intact females. We first waited for females to lay their first ootheca, which we collected, weighed and hatched (see next paragraph). Females were then randomly paired with new unrelated males, whose foretibial claws had been ablated in advance to prevent injury. Females that died prematurely before laying eggs (injured = 7, intact = 4) could not take part in this remating experiment. We also found that 9 females produced no offspring from the first ootheca, suggesting that no sperm was received from the first male, so these females were excluded from all subsequent analyses. Thus, the realised sample sizes for our remating experiment were: intact: *n* = 28, and injured: *n* = 26. Pairings lasted until females remated, cannibalised, or laid their second ootheca. We used the same mating-trial setup described above. Pairs were provided with blowflies inside their pairing enclosures at the same rate as normal feeding. We checked enclosures twice a day (morning and afternoon) for evidence of mating (the presence of a spermatophore), cannibalism (the absence of a male), and egg laying (the presence of an ootheca), and recorded the date and time, to the nearest hour, that these events were observed. Because sexual cannibalism in *M. caffra* is noncopulatory (Fea et al., 2013; Walker & Holwell, 2016), mating always proceeds without the male being eaten. However, there were some occasions where cannibalism occurred during a later interaction after mating or egg laying had occurred but before observations were made. We noted these instances for later consideration, but for this experiment we were primarily interested in the first outcome.

If injury alters female behaviour in a way that benefits injurious males, we predicted that injured females would take a longer time to remate compared to intact females, would be more likely to avoid remating entirely, would be more likely to cannibalise second males, or would be more likely to lay their second ootheca without remating. To assess the first prediction, we fitted a Gaussian linear model (LM) with latency to remate (time from the start of the trial to remating, in hrs) as the response variable, which was log transformed to meet the normality assumption. To test the second, third, and fourth predictions, we fitted three separate binomial GLMs in which remating outcome (1 = remating, 0 = no remating), cannibalism outcome (1 = cannibalism, 0 = no cannibalism), and ootheca outcome (1 = ootheca, 0 = no ootheca) were the respective binary response variables. In all four models, female treatment was the categorial fixed-effect predictor (intact vs. injured) with male and female age and body size fitted as the scaled and centred covariates. We allowed the treatment term to interact separately with each covariate term to account for the possibility that treatment effects were age-or size-dependent. Random effects for male and female family identity could not be included due to singularity issues.

### Terminal investment

To examine whether males benefit from injuring females by inducing a terminal investment response, we maintained females on a diet of 3 blow flies, fed twice a week, until they died. We collected and weighed all oothecae laid over the lifetime, with total ootheca weight used as a proxy for lifetime fecundity. We also counted the number of hatchlings that emerged from the first ootheca, and recorded the date of female death. The total number of females for which data were obtained for (a) survival, (b) ootheca weight, and (c) hatching counts was: (a) intact: *n* = 32, injured: *n* = 33, (b) intact: *n* = 28, injured: *n* = 26, and (c) intact: *n* = 32, injured: *n* = 33.

If injury causes females to invest terminally in a way that is adaptive for males, then we predicted that injured females would die faster after laying their first ootheca, and that these first oothecae would comprise a larger proportion of total lifetime fecundity in injured females compared to intact females. To test the first prediction, we fitted a mixed-effects Cox model in which the response variable was female age at death (time from injury procedure to death, in days) treated as a survival object. Observations were uncensored because deaths were recorded for all females. For the second prediction, we fitted a beta regression model with a logit link in which the response variable was the weight of the first ootheca divided by the weight of all oothecae, treated as a non-binomial proportion.

Because some proportions were unitary, we used the transformation 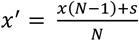 for beta-distributed data containing ones or zeroes, where *x* is the original proportion, *N* is the sample size, and *s* is a constant (0.5) (see Smithson & Verkuilen, 2006). In both models, female treatment was the categorical fixed-effect predictor which was allowed to interact separately with each of the scaled and centred numerical covariates of female age and female body size. Because some females had cannibalised a male or remated, we additionally included cannibalism status (i.e., whether cannibalism had previously occurred, 1 vs. 0) and remating status (i.e., whether a second mating had been achieved, 1 vs. 0) as binary predictors that were also allowed to interact with the treatment term to account for the possibility that treatment effects were dependent on these variables. Given the variable number of males that females encountered, male-based covariates were not included in the models. Female family identity was included as a random effect in the Cox model to account for relatedness between females, but was not included in the beta regression model due to singularity issues.

Because females should upregulate investment in early-produced offspring under terminal investment, we predicted that injured females would produce more offspring from the first ootheca compared to intact females. To test this prediction, we fitted a Poisson generalised linear mixed-effects model (GLMM) in which offspring count (number of hatchlings emerging from the first ootheca) was the Poisson response variable. The predictors and covariates were the same as those mentioned above for the beta regression model, except remating status and cannibalism status were not included, since females only remated and cannibalised after the first ootheca was laid. An observation-level random effect was added to the model to correct for overdispersion.

### General statistical methods

All analyses were performed in R version 4.4.1 (R Core Team, 2024). The binomial test was performed using the binom.test function (R Core Team, 2024). Mixed-effects models were fitted using the lmer and glmer functions in the lme4 package (Bates et al., 2015). The Cox model was fitted using the coxme function from the coxme package (Therneau, 2015), and the beta regression model was fitted using the betareg function from the betareg package (Cribari-Neto & Zeileis, 2010). All model covariates were scaled and centred to a mean of 0 and a standard deviation of 1 to ensure modelling on the same scale. A bobyqa optimiser was used to improve model convergence for the offspring GLMM. Singularity issues were dealt with by removing the offending random effects. To assess the significance of model variables, we used a model reduction approach in which the most nonsignificant terms were removed one at a time, starting with interaction terms, until a maximally reduced model was obtained that possessed only significant terms. Significance was determined by comparing models with and without the term of interest using likelihood ratio tests (LRTs) via the lrtest function from the lmtest package (Zeileis & Hothorn, 2002). Significant treatment-by-covariate interactions were assessed by comparing the marginal means of the linear trends in post hoc tests using the emtrends function from the emmeans package (Lenth, 2024). The treatment term was always retained in maximally reduced models regardless of its significance, with the intact group fixed as the reference level. For LRTs, we report Chi-square statistics (*𝒳*^2^) and p-values (*p*). For maximally reduced models, we report beta coefficients (*β*) and standard errors (*SE*) for significant predictors, including hazard ratios (*HR*) for the Cox model. For post hoc tests, we report the estimate (*est*.), standard error (*SE*), z ratio (*z*) and p-value (*p*). Female individual was the unit of replication. Deviations from our preregistered design and methodology are explained in Supplementary Material 1.

## RESULTS

### Female attractiveness

Injury did not alter the attractiveness of females (binomial test: 8 injured females chosen out of 16 trials, probability of success = 0.5, *p* = 1; Figure 1). However, chosen females that were injured were more likely to be older, and chosen females that were intact were more likely to be younger (mean ± SE age difference (intact – injured), in days: intact chosen: −7.38 ± 4.03; injured chosen: 8.75 ± 4.46; GLM: *β* = 0.25, *SE* = 0.15;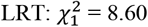, *p* = 0.003).

**Figure 1.**
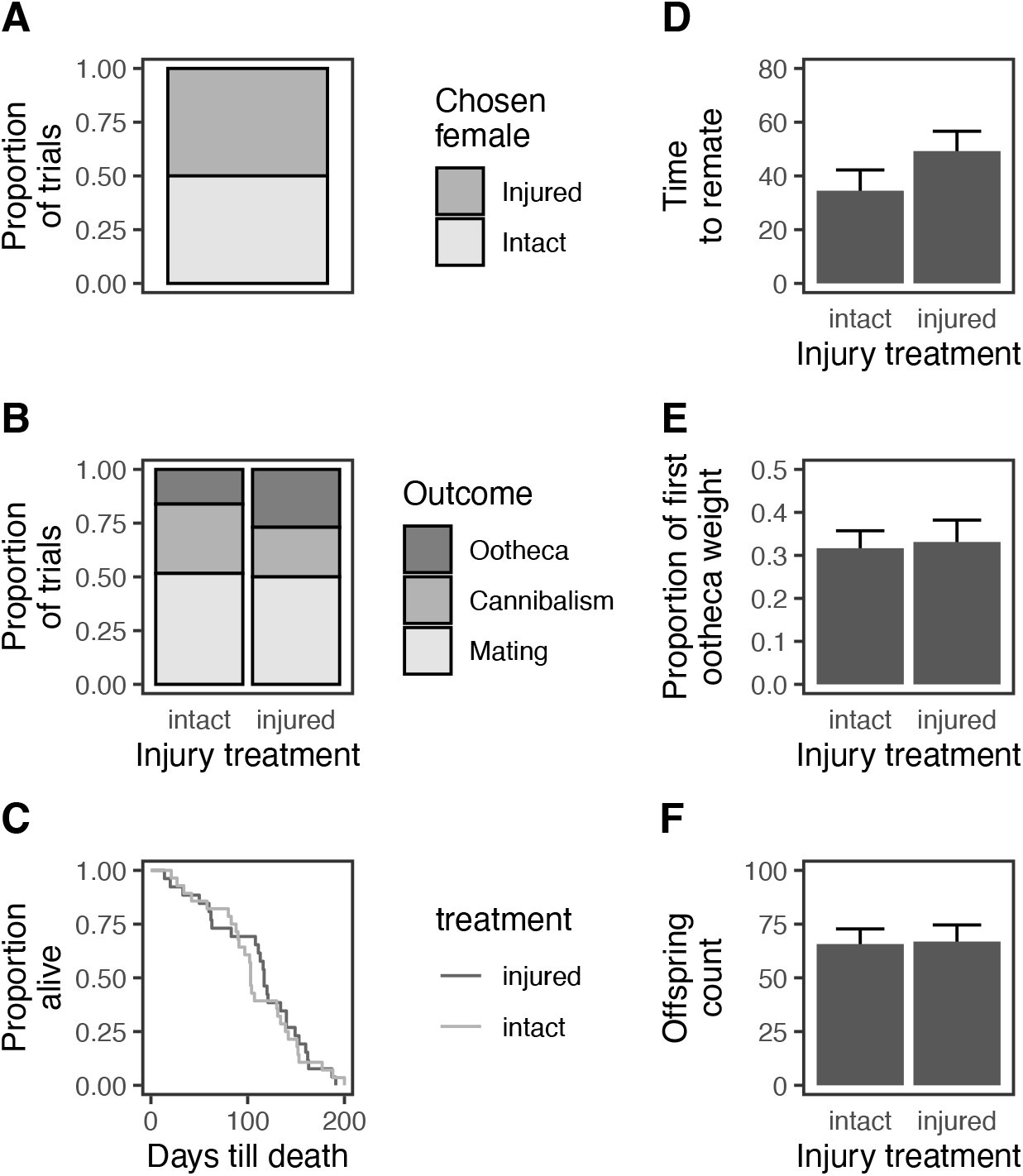
Plots showing the proportion of females chosen by males (A), the proportion of second mating trials ending in remating, cannibalism and egg laying (B), the proportion of females alive from first mating to death (C), time taken to remate, in hours (D), the proportional weight of the first ootheca out of the total weight of all oothecae (E), and the number of offspring emerging from the first ootheca (F). Error bars are means ± SE. The only significant difference between treatment groups is D.

### Remating and cannibalism

Injury had a significant effect on the latency to remate, with injured females taking 15 hours longer to remate than intact females (mean ± SE days to remate: intact: 34.50 ± 7.72, injured: 49.23 ± 7.41; GLM: *β* = 0.62, *SE* = 0.25;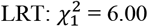, *p* = 0.01; Figure 1). Irrespective of the treatment group they belonged to, females additionally took longer to remate when paired with older males (GLM: *β* = 0.61, *SE* = 0.24;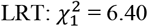, *p* = 0.01).

The effect of injury on the likelihood that females remated with a second male depended on the size of the male (GLM: *β* = 1.52, *SE* = 0.46;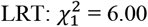, *p* = 0.01). Post hoc comparisons indicated that intact females were more likely to remate with smaller males (post hoc test: intact – injured: *est*. = −1.95, *SE* = 0.85, *z* = −2.29, *p* = 0.02). Regardless of treatment, females were also more likely to remate if the male was younger (GLM: *β* = −1.28, *SE* = 0.50;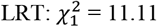, *p* = 0.001).

Injury did not affect the likelihood of cannibalism (GLM: *β* = −0.46, *SE* = 0.60;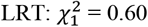, *p* = 0.44; Figure 1), or the likelihood to lay a second ootheca (GLM: *β* = 0.97, *SE* = 0.89;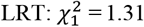, *p* = 0.25; Figure 1). Ovipositing instead of mating or cannibalising was more likely among females that were smaller (GLM: *β* = −1.40, *SE* = 0.56;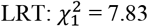, *p* = 0.005) or older (GLM: *β* = 0.88, *SE* = 0.38;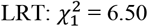, *p* = 0.01), and intact females were more likely to oviposit instead of remate or cannibalise when paired with larger males (GLM: *β* = 2.00, *SE* = 0.90;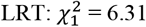, *p* = 0.01; post-hoc test: intact – injured: *est*. = 2.55, *SE* = 1.14, *z* = 2.23, *p* = 0.03).

### Terminal investment

Injury did not induce any change in female survival (Cox GLM: *β* = −0.08, *HR* = 0.92, *SE* = 0.33;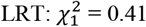, *p* = 0.52; Figure 1). However, females tended to die sooner if they were older at the time of mating i(Cox GLM: *β* = 0.30, *HR* = 1.35, *SE* = 0.17;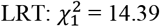, *p* = 0.01), or larger (Cox GLM: *β* = 0.61, *HR* = 1.84, *SE* = 0.23;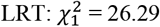, *p* < 0.001). Curiously, survival was enhanced among females that had previously cannibalised (Cox GLM: *β* = −0.82, *HR* = 0.44, *SE* = 0.20;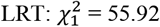, *p* < 0.001) and among females that had remated (Cox GLM: *β* = −0.63, *HR* = 0.53, *SE* = 0.18;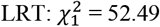, *p* < 0.001).

Injury did not induce females to terminally invest in reproduction, as the proportional contribution of the first ootheca to total fecundity in terms of weight did not differ between injured and intact females (GLM: *β* = −0.04, *SE* = 0.29;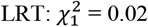, *p* = 0.90; Figure 1). However, we found evidence that female remating reduced the proportional contribution of the first ootheca weight to total ootheca weight (GLM: *β* = −0.35, *SE* = 0.15;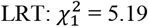, *p* = 0.02). Cannibalism following first oviposition had the same effect, reducing the weight of the first ootheca relative to the weight of all oothecae (GLM: *β* = −0.40, *SE* = 0.15;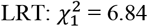, *p* = 0.009). Injury had no significant effect on the number of offspring hatching from the first ootheca (GLM: *β* = −0.29, *SE* = 0.46;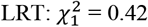, *p* = 0.52; Figure 1).

## DISCUSSION

We found little evidence that stabbing females in the abdomen is adaptive for *M. caffra* males. Injured females were not less attractive to other males, were not less likely to remate, did not avoid remating by cannibalising more, and were not more likely to oviposit a second time without remating. However, injury did cause females to delay remating, which could potentially provide a benefit to males if such a delay reduces sperm competition. In terms of reproduction, we observed no trade-off that could be interpreted as adaptive for males. Injured females did not upregulate investment in early-produced eggs, did not die sooner, and did not produce more offspring earlier. Taken together, our results suggest that sexual injury in this sexually cannibalistic species is likely to be a pleiotropic side-effect of selection acting on males to avoid being cannibalised.

The type of external stab wounds that *M. caffra* males inflict on females is expected to impose significant costs, including ruptured internal organs, infection, diverted resources for healing, and premature death (Lange et al., 2013). Indeed, we observed immediate and severe consequences of stabbing, with 17% of injured females dying prematurely without laying any eggs, indicative of a non-adaptive response. However, there did not appear to be any prolonged detrimental effects of being stabbed. Injured females that laid oothecae died at the same rate as intact females, and egg and offspring production did not differ between treatment groups, indicating no life-history trade-off for terminal investment. This suggests that females have robust physiological mechanisms to cope with sub-lethal injury, which is an expected outcome of rapid coevolution in response to male-inflicted harm (Arnqvist & Rowe, 2005). But harm itself is unlikely to be selected for in this species. Males only stab females if mounting attempts escalate to full-blown struggles, with such struggles enhancing mating success and reducing the likelihood of cannibalism (Burke & Holwell, 2021). Combined with our findings, this suggests that males are selected to wrestle aggressive females to subdue them for mating, with injury merely a side-effect of their anti-cannibalistic response.

A predicted outcome of adaptive injury is that wounded females should alter their remating behaviour to benefit the injurious male (Johnstone & Keller, 2000). In line with this prediction, we observed a significant 15-hour delay in the timing of remating among injured females. Such a delay could be beneficial in the wild if it reduces the likelihood that second males arrive to mate with injured females (for example, if the delay reflects lower female attractiveness). Because mantis density tends to be very low in wild populations (Barry et al., 2010; Maxwell, 1999), such an effect is feasible, but assessment of remating dynamics in the field would be necessary to confirm this. However, we suspect that this increase in the remating refractory period is unlikely to represent a benefit to males. First, the inter-oviposition period in this species is 1 to 2 weeks, so a 15-hour delay in remating is unlikely to be ecologically relevant. There was also no difference in the likelihood of remating between injured and intact females, indicating no real consequence of delaying remating. Second, most mated female pairs in our attractiveness experiment did not attract second males (34/50), and only 50% of both injured and intact females in our remating experiment actually remated, despite being paired continuously with males in small enclosures. This suggests that *M. caffra* females might become unattractive to males once mated, regardless of their injury status. Similar patterns have been observed in other mantises. For example, males of the Chinese mantis strongly prefer virgin over mated females in naturalistic settings (Lelito & Brown, 2008), and males of the false garden mantis also preferentially choose virgin females in the field, but freely mate with mated females in no-choice encounters (Barry et al., 2011). Male preference for virgin females is common in invertebrates (Richardson & Zuk, 2024), suggesting an important role for male mate choice (Bonduriansky, 2001). But females of some mantises also seem to actively avoid attracting second males. For example, females of the giant Asian mantis and European mantis stop calling to males pheromonally following a single copulation (Gemeno et al., 2005; Perez, 2005). If *M. caffra* females stop signalling in a similar way, then males may be unable to force mating reluctance beyond what females already naturally exhibit. Females might also benefit by halting production of sexual signals. Previous work indicates that mating a second time is suboptimal for *M. caffra* females (Burke & Holwell, 2023a), suggesting that already mated females might be selected to avoid attracting subsequent males so as to limit costs of mating multiple times. These females also receive enough sperm from a single mating to fertilise later-produced eggs (Burke & Holwell, 2023b), so a single mating may be sufficient to optimise female interests in this species.

An implicit assumption of adaptive manipulation of female remating through injury is that males must regularly be at risk of sperm competition. Such an assumption is likely to be met in *M. caffra* since both males and females are capable of mating multiply, and other polyandrous mantises show evidence of reduced paternity among first-mating males due to last male sperm precedence (Barry et al., 2011), which is generally common in insects (Simmons, 2002). Sperm competition also appears to be the driver of longer genital spines in the seed beetle *Callosobruchus maculatus* since the injuries that such spines inflict on the female reproductive tract facilitate greater absorption of seminal fluid and enhanced fertilisation success (Hotzy & Arnqvist, 2009). Such patterns suggest that sexual conflict over polyandry might generally favour injury as a side-effect of sperm competition (Hotzy & Arnqvist, 2009; Tong et al., 2021). But the link between mating system and injury evolution is unclear. Sexual conflict over monandry could potentially underlie harmful scape removal in orb-weavers if males monopolise fertilisations by injuring females (Mouginot et al., 2015), although costs of monandrous outcomes to female fitness have not been investigated in this system. Greater understanding of how injury is maintained in different mating systems could be a fruitful direction for future research on sexual injury.

To the best of our knowledge, our work here represents the first assessment of the adaptive harm hypothesis in a sexually cannibalistic species where males inflict injury with non-genitalic structures. We predicted that a unique way that males in sexually cannibalistic taxa could potentially manipulate female remating behaviour through injury could be by inducing more cannibalism toward later male suitors. Although injured females in our remating experiment were not more cannibalistic, investment in later-produced eggs increased among females that managed to cannibalise, suggesting that first males would benefit significantly if they could induce more aggression in their mates. The potential for sexually antagonistic manipulation of remating rates could be significant in sexually cannibalistic taxa, especially those where the consumption of males enhances female fecundity, as occurs in many mantises and spiders where sexual size dimorphism is not extreme (Wilder et al., 2009). Indeed, cannibalism appears to favour other forms of manipulation, such as deceptive pheromonal attraction of males by low quality females (Barry, 2015; Knapwerth & Burke, 2025). Male manipulation of female cannibalistic behaviour could also be widespread, mediated by external injury or via other mechanisms, such as seminal products or pheromones. However, we are unaware of any other studies that have investigated such manipulation in cannibalistic taxa.

We tried as much as possible to imitate real-life injuries when experimentally stabbing females, using a bespoke tool that ensured stab wounds were inflicted with exactly the same kind of weapon (a male claw), in the same location (ventral side of the female abdomen), and at the same depth (the length of a male claw). Nevertheless, it is possible that the violent wrestling that accompanies stabbing in this species (Burke & Holwell, 2021) inflicts greater injury than what we were able to achieve experimentally. Still, it is expected that female life-history should respond in a predictable way if injury is adaptive, regardless of the severity of harm (see Morrow et al., 2003). But, for the most part, we did not observe such patterns. It is also possible that natural injuries come with additional costs that we were unable to account for in our experiments. For example, claws of living males probably introduce contaminating agents to wounds, which could exacerbate infection and mortality. Understanding how injury affects male and female fitness in wild populations would be a valuable next step.

One of the requirements for experimentally testing hypotheses for adaptive versus pleiotropic harm is that the only factor that should be manipulated is the infliction of injury. For species with traumatic insemination, this is difficult to achieve. Experimenters have tried to prevent weaponised genitalia from inflicting injury internally by removing male genital spines with lasers (Hotzy et al., 2012) or by fixing wax to spines (Tong et al., 2021). The challenge with such an approach is that injury is not the only outcome that is manipulated: other functions of the injuring trait are also altered in the process. This makes it difficult to determine whether observed effects are due to the injuring organ no longer causing harm or due to it no longer operating normally, which could be quite problematic for primary sexual organs like genitalia. Indeed, phenotypic engineering experiments that limit genital spine functioning report lower mating success, reduced mating duration, and depressed fertilisation (Hotzy et al., 2012; Tong et al., 2021). Whether such effects are due to injury prevention or inhibition of genital function (or both) is unclear because the two factors cannot be readily disentangled. Extra-genital wounding—where injury is inflicted with structures other than male genitalia, such as spermatophores, teeth or claws (Lange et al., 2013)—permits greater control of the manipulation of injury *per se* without unintentional carry-over effects on mating outcomes. For example, our procedure of removing male claws is unlikely to have affected copulation duration, sperm transfer, or the ability of males to mate because such claws are primarily used for catching prey and are not directly used in fertilisation, allowing more robust assessment of the fitness effects of injury. Greater focus on the evolution of external wounding, such as bite wounds in spiders, mammals and fish (Chilvers et al., 2005; Pratt & Carrier, 2001; Sentenská et al., 2020) and spermatophore deposition in leaches and polychaetes (Lange et al., 2013) could be a fruitful direction for future research on the evolution of sexual harm.

## SUPPLEMENTARY MATERIAL

A registered report for this study can be found at: https://doi.org/10.17605/OSF.IO/Z5CE2. Some of the methods we had planned to utilise in our study as outlined in the registered report were later altered, either because we no longer considered them useful or because they could not be performed for statistical reasons. These alterations are listed below. First, we had initially intended to perform an analysis of the time that females took to lay their first ootheca. However, for terminal investment, we realised it was sufficient to assess differences in female survival and/or first ootheca weight and offspring count, without needing to also account for latency to lay, so we did not perform this analysis. Second, despite not being outlined in the original registered report, we conducted an additional analysis of latency to remate. The reason for this is that a number of previous studies have used remating interval as a metric to assess the adaptive harm hypothesis (e.g., Morrow et al., 2003), so we decided to include the same analysis for continuity. Third, although we did not mention it in the registered report, we also include an assessment of how male and female factors covary with male choice for the ‘female attractiveness’ data. This was done for the sake of completeness. Fourth, we tried to account for relatedness effects by including a random effects for male and female ootheca identity, but model singularity prevented this approach for most analyses. Fifth, we said in our registered report that we would include the timing of death relative to first oviposition (i.e., before vs. after) as a covariate in our survival analysis. We decided against this approach because of the low numbers involved. Instead, we report in the main results how many females of each treatment failed to oviposit. Finally, despite our modelling attempts, we could not account for the multiple use of certain males in the remating experiment because including male identity as a random effect generated model singularity.

## REFERENCES

Andrade, M. C. B. (1996). Sexual selection for male sacrifice in the Australian redback spider. Science, 271(5245), 70–72. 10.1126/science.271.5245.70

Arnqvist, G., & Rowe, L. (2002). Antagonistic coevolution between the sexes in a group of insects. Nature, 415, 787–789.

Arnqvist, G., & Rowe, L. (2005). Sexual Conflict. Princeton University Press.

Barry, K. L. (2015). Sexual deception in a cannibalistic mating system? Testing the Femme Fatale hypothesis. Proceedings of the Royal Society B: Biological Sciences, 282(1800), 20141428. 10.1098/rspb.2014.1428

Barry, K. L., Holwell, G. I., & Herberstein, M. E. (2008). Female praying mantids use sexual cannibalism as a foraging strategy to increase fecundity. Behavioral Ecology, 19(4), 710–715. 10.1093/beheco/arm156

Barry, K. L., Holwell, G. I., & Herberstein, M. E. (2009). Male mating behaviour reduces the risk of sexual cannibalism in an Australian praying mantid. Journal of Ethology, 27(3), 377–383. 10.1007/s10164-008-0130-z

Barry, K. L., Holwell, G. I., & Herberstein, M. E. (2010). Multimodal mate assessment by male praying mantids in a sexually cannibalistic mating system. Animal Behaviour, 79(5), 1165– 1172. 10.1016/j.anbehav.2010.02.025

Barry, K. L., Holwell, G. I., & Herberstein, M. E. (2011). A paternity advantage for speedy males? Sperm precedence patterns and female re-mating frequencies in a sexually cannibalistic praying mantid. Evolutionary Ecology, 25(1), 107–119. 10.1007/s10682-010-9384-3

Bates, D., Maechler, M., Bolker, B., & Walker, S. (2015). Fitting linear mixed-effects models using lme4. Journal of Statistical Software, 67(1), 1–48. 10.18637/jss.v067.i01

Blanckenhorn, W. U., Hosken, D. J., Martin, O. Y., Reim, C., Teuschl, Y., & Ward, P. I. (2002). The costs of copulating in the dung fly Sepsis cynipsea. Behavioral Ecology, 13(3), 353–358. 10.1093/beheco/13.3.353

Bonduriansky, R. (2001). The evolution of male mate choice in insects: a synthesis of ideas and evidence. Biological Reviews, 76(3), 305–339. 10.1017/S1464793101005693

Burke, N. W. (2024). Sexual cannibalism as a female resistance trait: a new hypothesis. Evolution, 78(4), 612–623. 10.1093/evolut/qpae017

Burke, N. W., & Holwell, G. I. (2021). Male coercion and female injury in a sexually cannibalistic mantis. Biology Letters, 16, 20200811. 10.1098/rsbl.2020.0811

Burke, N. W., & Holwell, G. I. (2023a). Costs and benefits of polyandry in a sexually cannibalistic mantis. 36(2), 412–423. 10.1111/jeb.14135

Burke, N. W., & Holwell, G. I. (2023b). Should females cannibalize with or without mating in the facultatively parthenogenetic springbok mantis? Animal Behaviour, 197, 113–121. 10.1016/j.anbehav.2023.01.007

Chapman, T. (2006). Evolutionary conflicts of interest between males and females. Current Biology, 16(17), 744–754. 10.1016/j.cub.2006.08.020

Chapman, T., Arnqvist, G., Bangham, J., & Rowe, L. (2003). Sexual conflict. Trends in Ecology & Evolution, 18(1), 41–47. 10.4161/fly.5.1.14459

Chapman, T., Liddle, L. F., Kalb, J. M., Wolfner, M. F., & Partridge, L. (1995). Cost of mating in Drosophila melanogaster females is mediated by male accessory-gland products. Nature, 373(6511), 241–244. 10.1038/373241a0

Chilvers, B. L., Robertson, B. C., Wilkinson, I. S., Duignan, P. J., & Gemmell, N. J. (2005). Male harassment of female New Zealand sea lions, Phocarctos hookeri: Mortality, injury, and harassment avoidance. Canadian Journal of Zoology, 83(5), 642–648. 10.1139/z05-048

Cribari-Neto, F., & Zeileis, A. (2010). Beta regression in R. Journal of Statistical Software, 34(2), 1– 24. 10.1201/9781003107347-10

Crudgington, H. S., & Siva-Jothy, M. T. (2000). Genital damage, kicking and early death. Nature, 407(6806), 855–856. 10.1038/35038154

Daly, M. (1978). The cost of mating. The American Naturalist, 112(986), 771–774. 10.1086/283319

Edvardsson, M., & Tregenza, T. (2005). Why do male Callosobruchus maculatus harm their mates? Behavioral Ecology, 16(4), 788–793. 10.1093/beheco/ari055

Elgar, M. A. (1992). Sexual cannibalism in spiders and other invertebrates. In M.A. Elgar & B. J. Crespi (Eds.), Cannibalism: Ecology and Evolution Among Diverse Taxa (pp. 128–155). Oxford University Press.

Fea, M. P., Stanley, M. C., & Holwell, G. I. (2013). Fatal attraction: sexually cannibalistic invaders attract naive native mantids. Biology Letters, 9(6), 1–4. 10.1098/rsbl.2013.0746

Fricke, C., Wigby, S., Hobbs, R., & Chapman, T. (2009). The benefits of male ejaculate sex peptide transfer in Drosophila melanogaster. Journal of Evolutionary Biology, 22(2), 275–286. 10.1111/j.1420-9101.2008.01638.x

Gavrilets, S. (2000). Rapid evolution of reproductive barriers driven by sexual conflict. Nature, 403(6772), 886–889. 10.1038/35002564

Gemeno, C., Claramunt, J., & Dasca, J. (2005). Nocturnal calling behavior in mantids. Journal of Insect Behavior, 18(3), 389–403. 10.1007/s10905-005-3698-y

Härdling, R., & Smith, H. G. (2005). Antagonistic coevolution under sexual conflict. Evolutionary Ecology, 19(2), 137–150. 10.1007/s10682-004-7917-3

Holland, B., & Rice, W. R. (1998). Perspective: chase-away sexual selection: antagonistic seduction versus resistance. Evolution, 52(1), 1–7. 10.2307/2410914

Hotzy, C., & Arnqvist, G. (2009). Sperm competition favors harmful males in seed beetles. Current Biology, 19(5), 404–407. 10.1016/j.cub.2009.01.045

Hotzy, C., Polak, M., Rönn, J. L., & Arnqvist, G. (2012). Phenotypic engineering unveils the function of genital morphology. Current Biology, 22(23), 2258–2261. 10.1016/j.cub.2012.10.009

Johnstone, R. A., & Keller, L. (2000). How males can gain by harming their mates: Sexual conflict, seminal toxins, and the cost of mating. The American Naturalist, 156(4), 368–377. 10.1086/303392

Kimura, K., & Chiba, S. (2015). The direct cost of traumatic secretion transfer in hermaphroditic land snails: Individuals stabbed with a love dart decrease lifetime fecundity. Proceedings of the Royal Society B: Biological Sciences, 282(1804), 20143063. 10.1098/rspb.2014.3063

Knapwerth, L., & Burke, N. W. (2025). Luring cannibal: dishonest sexual signalling in the springbok mantis. Functional Ecology, In product.

Lange, R., Reinhardt, K., Michiels, N. K., & Anthes, N. (2013). Functions, diversity, and evolution of traumatic mating. Biological Reviews, 88(3), 585–601. 10.1111/brv.12018

Lawrence, S. E. (1992). Sexual cannibalism in the praying mantid, Mantis religiosa: a field study. Animal Behaviour, 43(4), 569–583. 10.1016/S0003-3472(05)81017-6

Le Boeuf, B. J., & Mesnick, S. (1990). Sexual behavior of male northern elephant seals: i. lethal injuries to adult females. Behaviour, 116(1–2), 143–162.

Lelito, J. P., & Brown, W. D. (2006). Complicity or conflict over sexual cannibalism? Male risk taking in the praying mantis Tenodera aridifolia sinensis. American Naturalist, 168(2), 263–269. 10.1086/505757

Lelito, J. P., & Brown, W. D. (2008). Mate attraction by females in a sexually cannibalistic praying mantis. Behavioral Ecology and Sociobiology, 63(2), 313–320. 10.1007/s00265-008-0663-8

Lenth, R. V. (2024). emmeans: estimated marginal means, aka least-squares means. R package version 1.10.4. https://CRAN.R-project.org/package=emmeans.

Lessells, C. M. (2005). Why are males bad for females? Models for the evolution of damaging male mating behavior. American Naturalist, 165, S46–S63. 10.1086/429356

Maxwell, M. R. (1999). Mating behaviour. In F. R. Prete (Ed.), The Praying Mantids (pp. 69–89). Johns Hopkins University Press.

Maxwell, M. R., Barry, K. L., & Johns, P. M. (2010). Examinations of female pheromone use in two praying mantids, Stagmomantis limbata and Tenodera aridifolia sinensis (Mantodea: Mantidae). Annals of the Entomological Society of America, 103(1), 120–127. 10.1603/008.103.0115

Michiels, N. K. (1998). Mating conflicts and sperm competition in simultaneous hermaphrodites. In T.R. Birkhead & A.P. Møller (Eds.), Sperm Competition and Sexual Selection (pp. 219–254). Academic Press.

Moore, A. J., Gowaty, P. A., & Moore, P. J. (2003). Females avoid manipulative males and live longer. Journal of Evolutionary Biology, 16(3), 523–530. 10.1046/j.1420-9101.2003.00527.x

Morrow, E. H., & Arnqvist, G. (2003). Costly traumatic insemination and a female counter-adaptation in bed bugs. Proceedings of the Royal Society B: Biological Sciences, 270(1531), 2377–2381. 10.1098/rspb.2003.2514

Morrow, E. H., Arnqvist, G., & Pitnick, S. (2003). Adaptation versus pleiotropy: Why do males harm their mates? Behavioral Ecology, 14(6), 802–806. 10.1093/beheco/arg073

Mouginot, P., Prügel, J., Thom, U., Steinhoff, P. O. M., Kupryjanowicz, J., & Uhl, G. (2015). Securing paternity by mutilating female genitalia in spiders. Current Biology, 25(22), 2980– 2984. 10.1016/j.cub.2015.09.074

Mouginot, P., & Uhl, G. (2019). Females of a cannibalistic spider control mutilation of their genitalia by males. Behavioral Ecology, 30(6), 1624–1631. 10.1093/beheco/arz127

Nakata, K. (2016). Female genital mutilation and monandry in an orb-web spider. Biology Letters, 12(2). 10.1098/rsbl.2015.0912

Parker, G. A. (1979). Sexual selection and sexual conflict. In N. Blum & M. Blum (Eds.), Sexual Selection and Reproductive Competition in Insects (pp. 123–166). Academic Press. 10.1016/B978-0-12-108750-0.50010-0

Perez, B. (2005). Calling behaviour in the female praying mantis, Hierodula patellifera. Physiological Entomology, 30(1), 42–47. 10.1111/j.0307-6962.2005.00426.x

Perry, J. C., & Rowe, L. (2015). The evolution of sexually antagonistic phenotypes. Cold Spring Harbor Perspectives in Biology, 7(6), 1–18. 10.1101/cshperspect.a017558

Pollo, P., Burke, N. W., & Holwell, G. I. (2021). Effects of male and female personality on sexual cannibalism in the Springbok mantis. Animal Behaviour, 182, 1–7.

Pratt, H. L., & Carrier, J. C. (2001). A review of elasmobranch reproductive behavior with a case study on the nurse shark, Ginglymostoma cirratum. Environmental Biology of Fishes, 60, 157– 188.

R Core Team. (2024). R: A language and environment for statistical computing (4.4.1). R Foundation for Statistical Computing. https://www.r-project.org/

Richardson, J., & Zuk, M. (2024). Meta-analytical evidence that males prefer virgin females. Ecology Letters, 27(1), e14341. 10.1111/ele.14341

Rowe, L., Cameron, E., & Day, T. (2005). Escalation, retreat, and female indifference as alternative outcomes of sexually antagonistic coevolution. American Naturalist, 165(S5), S5–S18. 10.1086/429395

Schneider, J. M. (2014). Sexual cannibalism as a manifestation of sexual conflict. Cold Spring Harbor Perspectives in Biology, 6(11), 1–16. 10.1101/cshperspect.a017731

Sentenská, L., Šedo, O., & Pekár, S. (2020). Biting and binding: an exclusive coercive mating strategy of males in a philodromid spider. Animal Behaviour, 168, 59–68. 10.1016/j.anbehav.2020.08.001

Simmons, L. W. (2002). Sperm Competition and its Evolutionary Consequences in the Insects. Princeton University Press.

Smithson, M., & Verkuilen, J. (2006). A better lemon squeezer? Maximum-likelihood regression with beta-distributed dependent variables. Psychological Methods, 11(1), 54–71. 10.1037/1082-989X.11.1.54

Sontag, C. (1971). Spectral sensitivity studies on the visual system of the praying mantis, Tenodera sinensis. Journal of General Physiology, 57(1), 93–112. 10.1085/jgp.57.1.93

Therneau, T. M. (2015). coxme: mixed effects Cox models (R package version 2.2-5). R package version 2.2-5. https://cran.r-project.org/package=coxme

Tong, X., Wang, P. Y., Jia, M. Z., Thornhill, R., & Hua, B. Z. (2021). Traumatic mating increases anchorage of mating male and reduces female remating duration and fecundity in a scorpionfly species. Proceedings of the Royal Society B: Biological Sciences, 288(1952). 10.1098/rspb.2021.0235

Walker, L. A., & Holwell, G. I. (2016). Sexual cannibalism in a facultative parthenogen: The springbok mantis (Miomantis caffra). Behavioral Ecology, 27(3), 851–856. 10.1093/beheco/arv221

Wilder, S. M., Rypstra, A. L., & Elgar, M. A. (2009). The importance of ecological and phylogenetic conditions for the occurrence and frequency of sexual cannibalism. Annual Review of Ecology, Evolution, and Systematics, 40(1), 21–39. 10.1146/annurev.ecolsys.110308.120238

Zeileis, A., & Hothorn, T. (2002). Diagnostic checking in regression relationships. R News, 2, 7–10.

